# Improving antibody thermostability based on statistical analysis of sequence and structural consensus data

**DOI:** 10.1101/2021.01.28.428721

**Authors:** Lei Jia, Mani Jain, Yaxiong Sun

## Abstract

The use of Monoclonal Antibodies (MAbs) as therapeutics has been increasing over the past 30 years due to their high specificity and strong affinity towards the target. One of the major challenges towards their use as drugs is their low thermostability, which impacts both efficacy as well as manufacturing and delivery. To aid the design of thermally more stable mutants, consensus sequence-based method has been widely used. These methods typically have a success rate of about 50% with maximum melting temperature increment ranging from 10 to 32°C. In order to improve the prediction performance, we have developed a new and fast MAbs specific method by adding a 3D structural layer to the consensus sequence method. This is done by analyzing the close-by residue pairs which are conserved in more than eight hundred MAbs’ 3D structures. Major advantage of this structural level assessment is in significantly reducing the false positives by almost half from the consensus sequence method alone. In summary, combining consensus sequence and structural residue pair covariance methods, we developed an inhouse application for predicting human MAb thermostability to guide protein engineers to design more stable molecules. This application has shown success in designing MAb engineering panels in multiple biologics programs.

## Introduction

Monoclonal Antibodies (MAbs) have become one of the most important classes of therapeutics in various disease areas. Half of the top 10 bestselling drugs in year 2019 are MAbs [1]. Thermostability is a basic biophysical property of MAbs. But it can be a major concern in the development of protein therapeutics due to its impact on both efficacy as well as manufacturing and delivery [2]. Low thermostability can cause MAbs to denature and aggregate [3] which can further lead to loss in binding potency [4], lower purity in manufacturing [5], and shortened shelf life [6]. Optimizing MAbs’ thermostability attribute is among the fundamental protein-engineering processes for a therapeutic MAb discovery project [7–11]. A high-throughput and accurate in silico prediction method can be helpful to eliminate liable molecules as early as possible and design mutations to improve thermostability of lead molecules.

There are multiple approaches to predict protein thermostability and engineer protein to improve its stability [12, 13]. Protein consensus sequence-based methodology has been applied to improve protein thermostability for over 20 years [14–18]. In general, the success rate for this statistics-based method is about 50% with maximum melting temperature increase ranging between 10 and 32°C [19]. Since MAbs all fold into conserved structures [20], we can push the consensus method to structural level and study the covariance relationship between residue pairs on 3D protein structures. Structural level assessment helped to significantly decrease false positives from the consensus sequence method alone.

Combining consensus sequence and structural residue pair covariance methods, we developed an inhouse application for predicting human MAb thermostability to guide protein engineers to design more stable molecules. The consensus method is trained by ~25K and ~12K human heavy and light chain variable region sequences, respectively, from The international ImMunoGeneTics information system (IMGT) http://www.imgt.org/ [21]. The structural covariance method is trained by over 800 curated high-resolution crystal structures, and a scoring system has been developed to evaluate pairwise residue interaction with confidence in consideration. The application, which consists of about 1500 lines of python codes, has been incorporated in our antibody engineering workflow. It has shown success in designing MAb engineering panels in multiple biologics programs. Further development areas include enriching the training data for human MAb prediction (improve accuracy based on statistical significance), developing predictive models for other species’ antibodies, e.g. camelid heavy chain only antibody, and seeking applications for multi-specific antibody engineering.

## Methods

### Sequence Based Consensus Scoring

MAb sequences from IMGT were used as the starting point. We specifically focused on the variable domains of human MAbs. From IMGT (as of year 2016), we obtained 35,614 human VH, 7,674 human VK, and 5,430 human VL sequences. The human germline sequence definition was taken from the V BASE, https://www2.mrc-lmb.cam.ac.uk/vbase/. All antibody sequences from IMGT were assigned to a germline type and germline family based on sequence alignment to the germline sequences. In order to have a cleaner germline annotation, for each sequence being used for consensus calculation, we set up 80% sequence similarity threshold as a filter to construct the consensus sequence data base. The filtering process yields 25,220 VH, 7,190 VK, and 4,789 VL sequences. Table 1 shows the number of sequences in each germline family.

**Table 1:** Number of sequences in each germline family.

| Germline Family | No. of Sequences | Germline Family | No. of Sequences | Germline Family | No. of Sequences |
| --- | --- | --- | --- | --- | --- |
| VH1 | 3111 | VK1 | 3412 | VL1 | 1307 |
| VH2 | 619 | VK2 | 1049 | VL2 | 1039 |
| VH3 | 15610 | VK3 | 2143 | VL3 | 1465 |
| VH4 | 3490 | VK4 | 516 | VL4 | 154 |
| VH5 | 1005 | VK5 | 29 | VL5 | 110 |
| VH6 | 1231 | VK6 | 41 | VL6 | 300 |
| VH7 | 154 |  |  | VL7 | 132 |
|  |  |  |  | VL8 | 206 |
|  |  |  |  | VL9 | 38 |
|  |  |  |  | VL10 | 38 |

The consensus sequences were calculated based on three different levels: 1, all sequences in VH, VK, or VL sequence database; 2, all sequences at the germline family level; 3, all sequences at the germline level. To calculate the consensus sequence, the sequences were annotated and aligned following an Amgen in house numbering scheme, which is similar to the AHo numbering scheme. At each residue position, the amino acid with the highest frequency is defined as consensus amino acid.

The idea behind the consensus sequence-based protein stability engineering method is that “a conserved residue is more likely to be stabilizing than a random mutation at that same position”. For a given amino acid in the query sequence (the sequence that needs to be evaluated for thermostability) we define the consensus score as a free energy change (ΔΔG_AA_) by using Boltzmann distribution theory:

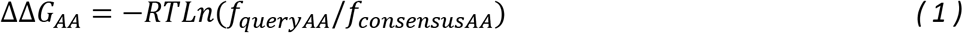

Where, the R is the Boltzmann constant, T is temperature (298K), f_queryAA_ is the frequency of the query amino acid at a given position, f_consensusAA_ is the frequency of the consensus amino acid at that same position. ΔΔG_AA_ measures the effect of a single residue change to consensus amino acid on the stability of the antibody molecule and, a positive ΔΔG_AA_ reflects increased stability. The consensus score for the whole antibody sequence (ΔΔG_Seq_) is the sum of the consensus score of each individual amino acid’s consensus score as shown in equation 2. ΔΔG_Seq_ measures the overall effect of all the single amino acid changes to consensus residues on the stability of the MAb and the higher ΔΔG_Seq_ is the more stable the MAb will be.

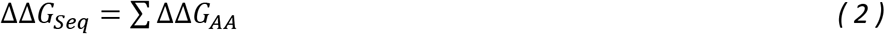

Applying the consensus analysis can be based on the whole IMGT database level, germline family level or germline level, ranging from high level to low level sequence coverage. The higher, the level, e.g at IMGT database level, the more sequences are available in the gene pool, and thus more statistically meaningful results can be obtained. On the other hand, the higher, the level, the sequences used to calculate consensus are less specific to the query sequence (lower sequence homology). Thus, the results may not be accurate. The scope of the analysis can be based on Complementarity-determining region (CDR), framework region (FR), whole Fv or the whole Fab region. The consensus method works best for the FR region as FR region provides structural stability to the MAb. For the purpose of this study we have focused on the germline family level (the sequence homology and statistical confidence are appropriate at this level), FR region (provides structural stability) and single point mutation (provides cleaner validation). Hence, in our results section the ΔΔG_Seq_ (ddG) values reflect the effect of mutating one position in the query sequence to consensus or germline residue.

### Metric to select the best ddG cutoff

ddG cutoff serves as our confidence threshold for the predicted thermostabilizing mutations. If for a query sequence, the ddG score is higher than the ddG cutoff value we suggest the protein engineers with high confidence that the mutation to consensus residue will be thermostabilizing and will help in improving the melting temperature. The predictions from the consensus sequence method can fall under four categories as shown in Figure 1. True Positives (TP) (mutations predicted to have ddG values greater than or equal to the ddG cutoff and experimentally observed melting temperature (T_m_) difference between the mutant and the query sequence is greater than or equal to the dT_m_ cutoff), False Positives (FP) (mutations predicted to have ddG values greater than or equal to the ddG cutoff but experimentally observed melting temperature difference between the mutant and the query sequence is less than the dT_m_ cutoff), True Negatives (TN) (mutations predicted to have ddG values less than the ddG cutoff and experimentally observed melting temperature difference between the mutant and the query sequence is lower than the dT_m_ cutoff) and False Negatives (FN) (mutations predicted to have ddG values less than the ddG cutoff but experimentally observed melting temperature difference between the mutant and the query sequence is greater than or equal to the dT_m_ cutoff).

**Figure 1:**
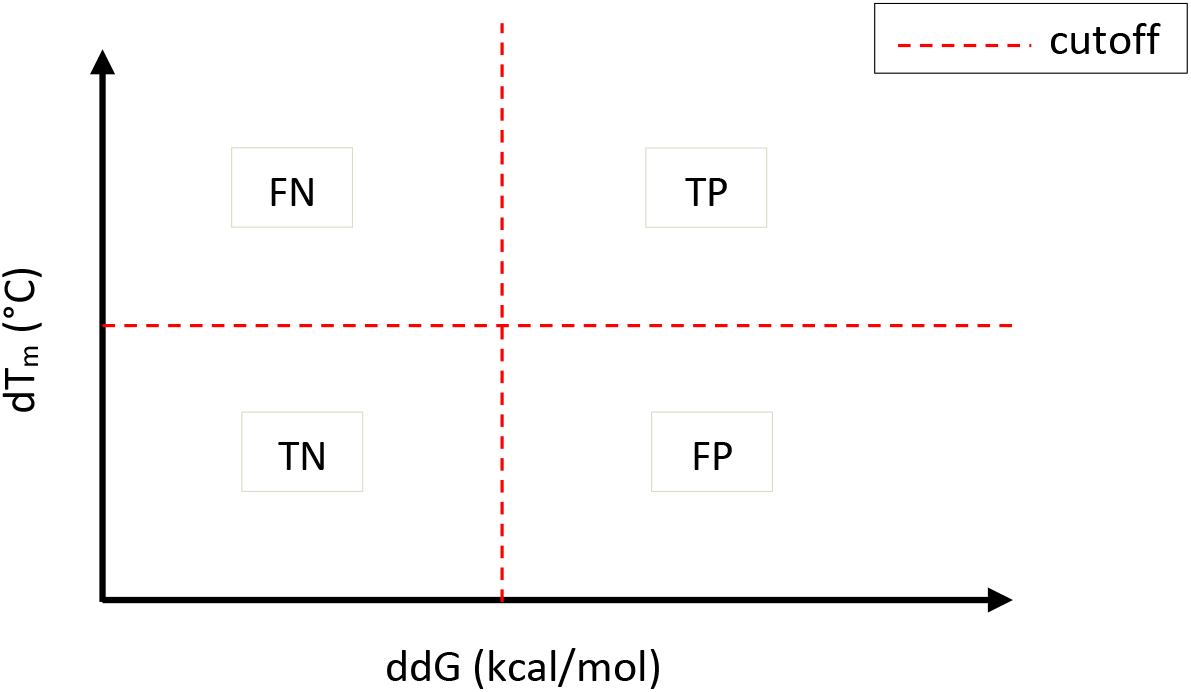
The x-y plot is showing the four categories into which the predictions from the consensus sequence method can fall. The x-axis represents the ddG values predicted by consensus sequence method and y-axis is showing the experimentally observed melting temperature difference between the mutant and reference molecule. The red dotted lines represent the cutoff values for ddG and dT_m_.

The dT_m_ cutoff is dependent on the experimental error in T_m_ measurements. The idea behind the metric to select the best ddG cutoff is based on maximizing TP, minimizing FP and at the same time maximizing value for true predictions (TP+TN). We care about TP the most, and then reducing FP and then maximizing our total true predictions. We devised and explored four different metrics along with Precision as shown in Table 2.

**Table 2:** Different metrics explored for selecting the best ddG cutoff. Note: Criteria is based on navigation from lowest to highest ddG cutoff values.

| Metric | Formulae | Criteria for choosing ddG cutoff <sup>note</sup> |
| --- | --- | --- |
| Precision | $TP/(TP+FP)$ | Not defined |
| Metric 1 | $TP*(TN-FP)/(FP*TN)$ | >0 |
| Metric 2 | $TP*(TN-FP)$ | $\geq 0$ |
| Primary Metric (P Metric) | $TP+TP*(TN-FP)$ | >0 |
| Fine-tuning Metric (F Metric) | $(TP-TN)/(TN-FP)$ | $\geq 0$ |

Finally, we decided on the using two metrics Primary Metric (P Metric) and Fine-tune metric (F Metric) in conjunction with one-another. The first step is to titrate the ddG cutoff values and sort them, we used a step size of 0.1. The criteria for choosing the best ddG cutoff is defined as we navigate from lowest to highest ddG cutoff. The selection of best ddG cutoff is defined with respect to the ddG value at which P metric > 0 (ddG_cutoff(i)_) and if the value of F metric is < 0 at ddG_cutoff(i)_ then we will pick a ddG cutoff one before the one which has P metric > 0 i.e. ddG_cutoff(i-1)_. Otherwise, if F metric is positive at ddG_cutoff(i)_ we will pick that as the best ddG cutoff. This procedure is described in the following pseudocode. The key point is to traverse possible ddG cutoff values from lowest to highest and finer titrations of ddG cutoff gives us a better cutoff value.

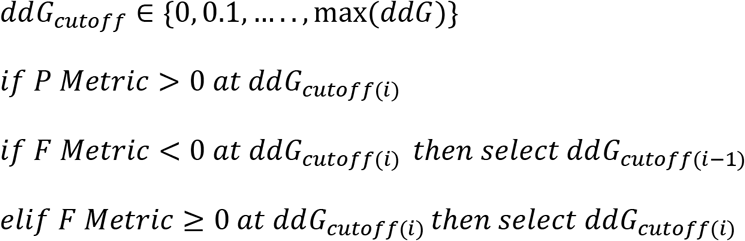

### Consensus structure-based MAb residue-pair covariance analysis

We obtained 841 crystal structures of human antibody variable domain from IMGT 3D structure database. Pairwise residue distance matrix was calculated for each of Fv structure using only FR residues. We used the following two criteria to flag a pair of residues as close-by: 1. only calculate amino acid residue pairs which are separated by more than 2 amino acids (to avoid analyzing adjacent pairs and loop tip at the beta-hairpin); 2. pick minimum distance between all side chain heavy atoms of the amino acid pairs, used 4 Å distance as cutoff. Residue pairs which have minimum distance between any side chain heavy atom less than 4 Å are flagged as close-by residue pairs. The residue numbers on the variable domain are uniform for all 841

MAbs based on Amgen in house numbering system. After all close-by residue pairs were calculated for all 841 antibody structures, we used 100 occurrences as a cutoff to mark consensus close-by residue pairs. This yielded 257 close-by residue pairs and we used these for assessing the residue synergy in the probe antibody.

Close-by residue pairs are preferred if the 2 residues have opposite charge leading to columbic interaction, or both having high hydrophobicity for favored van der Waals interaction (packing), or both are aromatic for favored stacking interaction. In order to quantify the favorable interactions, we developed a scoring system:

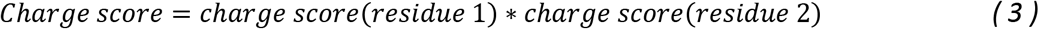

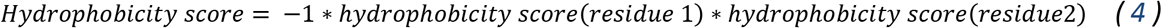

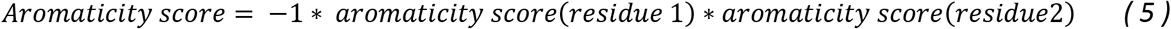

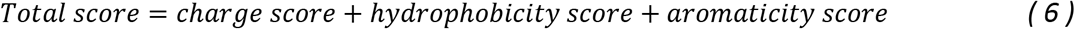

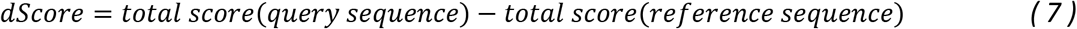

The charge and hydrophobicity scores are defined by amino acid charge and hydrophobicity in references [22, 23]. The aromaticity score is based in reference [23] and defined with our empirical adjustment as following (tryptophan: 1, phenylalanine and tyrosine: 0.8, histidine: 0.3, arginine: 0.3, and the rest of amino acids are 0). The reference sequence can either be the germline sequence or the consensus sequence. Similar to energy, the lower the dScore, the more favorable are the residue pairs and thus more favorable is the antibody which contain such residue pairs. One residue can be present in multiple close-by residue pairs. For protein engineering purpose, each residue’s impact on protein stability is summed up by their contribution to every close-by residue pairs which contains it, with adjustment of a confidence level:

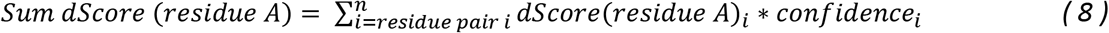

Confidence is defined as the ratio of number of times a pair of residues is present as close-by residue pairs to the total 841 structures:

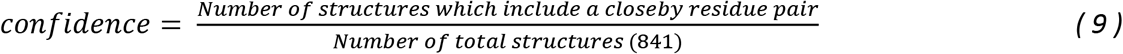

The final output of the MAb residual covariance analysis is the sum dScore (equation 8). Comparing to the reference MAb, if this score is a positive number, it means that the residue is less stable than the residue in the reference MAb. A mutation to consensus residue is suggested to stabilize the query MAb.

This method combining consensus sequence method with consensus structure-based MAb residue-pair covariance analysis is implemented in Python programming language. A graphical user interface is also developed with Pipeline Pilot for deployment. It requires as input Fv domain sequence of MAbs in fasta format (annotated and aligned following the Amgen numbering scheme). The workflow of the inhouse Pipeline Pilot tool includes two steps, first step is to calculate the consensus sequence and the ddG score (equation 1) for all residues in query sequence in the FR region and; second step is to perform the structural filtering. The tool outputs a csv file with thermostabilizing mutation suggestions for each MAb and includes the ddG score. The users have an optional flag to turn off structural filtering as well i.e. retrieve mutation suggestions based only on the consensus sequence method. By default, structural filtering is turned on and the output only includes thermostabilizing mutations which have passed structural filtering.

## Results

### Evaluate the consensus sequence method with published data

To evaluate the consensus sequence prediction method, we used a stability engineering study of the single chain fragment variable domain (scFv) of an antibody published by Monsellier et al. [24] In that study the authors used consensus method to predict point mutations of scFv to improve thermostability. Experimental free energy change (ddG) was reported to assess the computational method. Table 3 shows the experimental ddG, calculated ddG from that paper and from our method for the mutants. Our calculated ddG values are similar to those reported in the paper and both calculated ddG values per variant are in the same positive direction as the experimental ddG, which indicates good prediction outcome. Through this comparison, we validated the consensus sequence method that we have implemented.

**Table 3:** Experimental and calculated ddG (in kcal/mol) to validate our consensus sequence method.

| Variants | Mutations | Experimental ddG | Calculated ddG from Monsellier et al. | Calculated ddG from our work |
| --- | --- | --- | --- | --- |
| L1 | L-Q40P,L-K42Q | 0.6 | 3.1 | 4.2 |
| L2 | L-Q45K | 4.1 | 1.3 | 0.9 |
| L3 | L-K74T | 2.9 | 0.7 | 1.1 |
| L4 | L-N76S | 0.6 | 1.5 | 1.7 |
| L5 | L-G84A, L-S85T | 2.5 | 2.3 | 3.5 |
| H1 | H-S15G | 0.1 | 1 | 0.7 |
| H2 | H-S61E, H-A62K, H-L63F | 1.6 | 2.1 | 2.5 |
| H3 | H-H83T, H-T84S, H-D85E | 0.9 | 5.9 | 6.7 |

### Selecting the optimal ddG cutoff for thermostability prediction and MAb engineering

We noted that the consensus sequence stability method does not have high accuracy to confidently predict the actual melting temperature (T_m_) and experimental ddG directly. Protein engineers desire to have a high throughput prediction method to predict variants’ stability and propose what mutations they can make to improve stability. So, we further developed this method to be a classification tool to meet the above protein engineers need. For classification, we need to determine a ddG cutoff value to confidently predict the amino acid stability at a given position. identify the amino acid position which can be engineered to improve stability. We focused on the FR in the MAb Fv and analyzed only single point mutations for cleaner validation. We used a set of 234 MAb thermostability data in T_m_ representing 201 single point mutation pairs to determine the optimal ddG cutoff. The dT_m_ cutoff used in our work is 0.5 °C, it’s chosen based on the distribution of dT_m_ measurements in our experimental data as shown in Figure 2 and suggestion from the analytical scientist who performed the measurements (based on the experimental error bar in T_m_ measurements by Differential Scanning Fluorimetry (DSF)). We devised four metrics namely Metric 1, Metric 2, P Metric and F metric as shown in Table 2. We compared the TP, FP, TN and FN statistics for different ddG cutoff values using these four metrics and the precision as shown in Table S1. The criteria for selecting the optimal ddG cutoff using each of these five metrics is shown in Table 2. For the precision we were not able to come up with any definite criteria based on which we can choose the optimal ddG cutoff.

**Figure 2:**
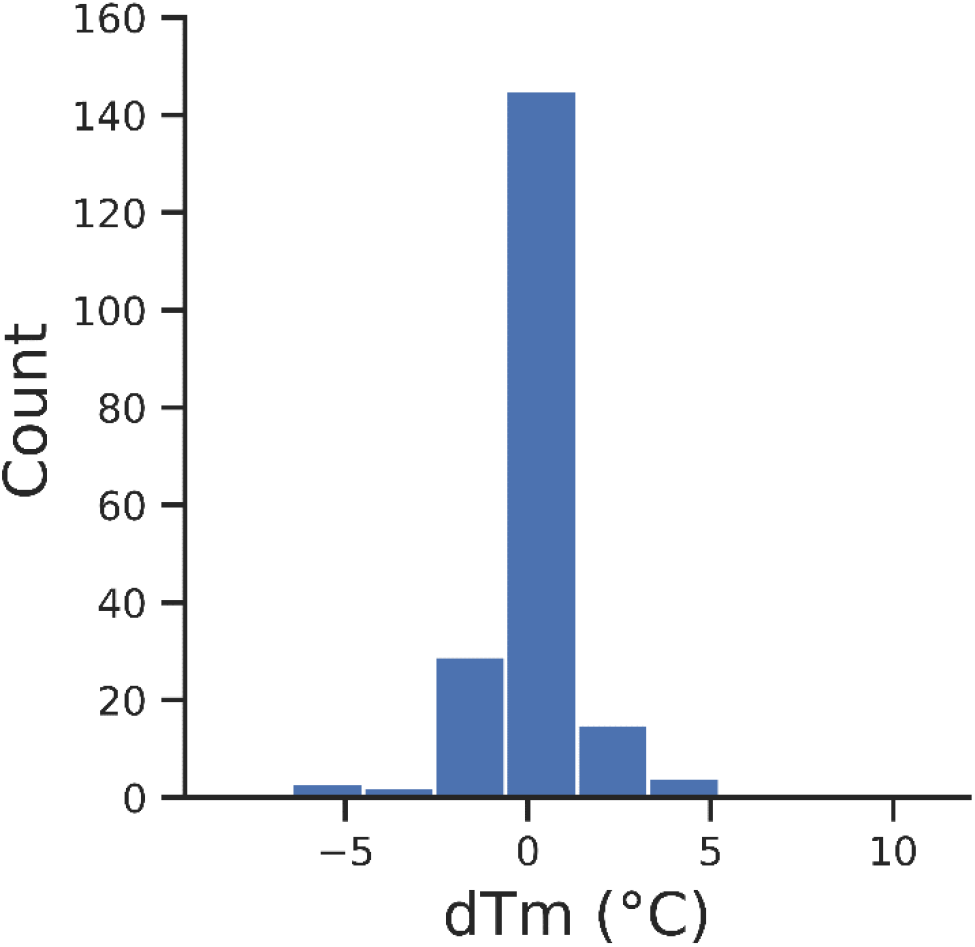
The frequency distribution of dT_m_ for 201 single point mutation pairs in our dataset.

If we navigate from the lowest to the highest ddG cutoff values at the fixed dT_m_ cutoff of 0.5 °C we observed that TP and FP will reduce but TN and FN will increase as shown in Table S1. Since we care the most about maximizing TP, we start the navigation from the lowest ddG cutoff as TP will be the highest. The second goal is to minimize FP as much as possible without sacrificing many TP. Based on this rationale, we chose P Metric and F Metric together as defined in the methods section, for further information refer to the analysis shown in Table S1 and Figure S1. Based on our P Metric and F Metric we selected 1.7 kcal/mol as the optimal ddG cutoff for the consensus sequence method. The selected optimal ddG cutoff depends on the data set and prediction confidence goal. So, in another use case, this value can be different from the value we obtained. In the next section, we demonstrated that this cutoff value is further optimized based on another use case (adding consensus structural filter). In future, with more data, this cutoff can be further optimized.

### Consensus structure-based MAb residue-pair analysis can significantly decrease the false positive prediction rate

Human MAb has a conserved structural fold. Described by IMGT’s Colliers de Perles illustration, the variable domain of heavy and light chains each has 9 anti-parallel beta-sheets, which form the FR of the variable domain. 3 out of 4 loops in connecting the beta-sheets build up the CDR region. The consensus based on a set of MAb crystal structures can provide a general representation of their overall structural features. This is the rationale behind the consensus structure-based MAb residue-pair method. This method is high throughput but not highly accurate to predict the stability directly. So, we only use this method as a filter on top of the consensus sequence method. We demonstrated that this structural method can significantly decrease the FP of the consensus sequence method prediction.

We used the same set of 234 MAb thermostability data in T_m_ representing 201 single point mutation pairs for validating the consensus structure-based MAb residue-pair method. 154 single point mutation pairs out of 201 passed the structural filter i.e. they had a positive Sum dScore as described in equation 8. For practical application purposes, we optimized the ddG cutoff again using these 154 data points as the protein engineers will only be looking at the mutations which passed the structural filter. By applying P Metric and F Metric on these 154 single point mutation pairs at the fixed dT_m_ cutoff of 0.5 °C, we obtained the optimal ddG cutoff of 1.3 kcal/mol (for further details refer to the Table S2 and Figure S2). For our further analysis, we have used this practical optimal ddG cutoff of 1.3 kcal/mol.

Figure 3 is a scatterplot of dT_m_ (difference of experimental T_m_ between the mutant and the parental molecules) VS ddG (predicted from the consensus sequence method) for all 201 single point mutation pairs. We also computed confusion matrix and precision comparing the performance between without and with structural filtering (Table 4). Since we care most about maximizing TP and minimizing FP, we would aim to have as high precision as possible. We can observe that structural filtering leads to precision improvement. To visualize the effect of consensus structure-based MAb residue-pair method in reducing FP, we plotted a bar plot as shown in Figure 4. We can see structural filtering helped in significantly reducing the FP without sacrificing much of the TP. We observed a consistent effect of structural filtering on reducing number of FP at different ddG cutoffs also.

**Figure 3:**
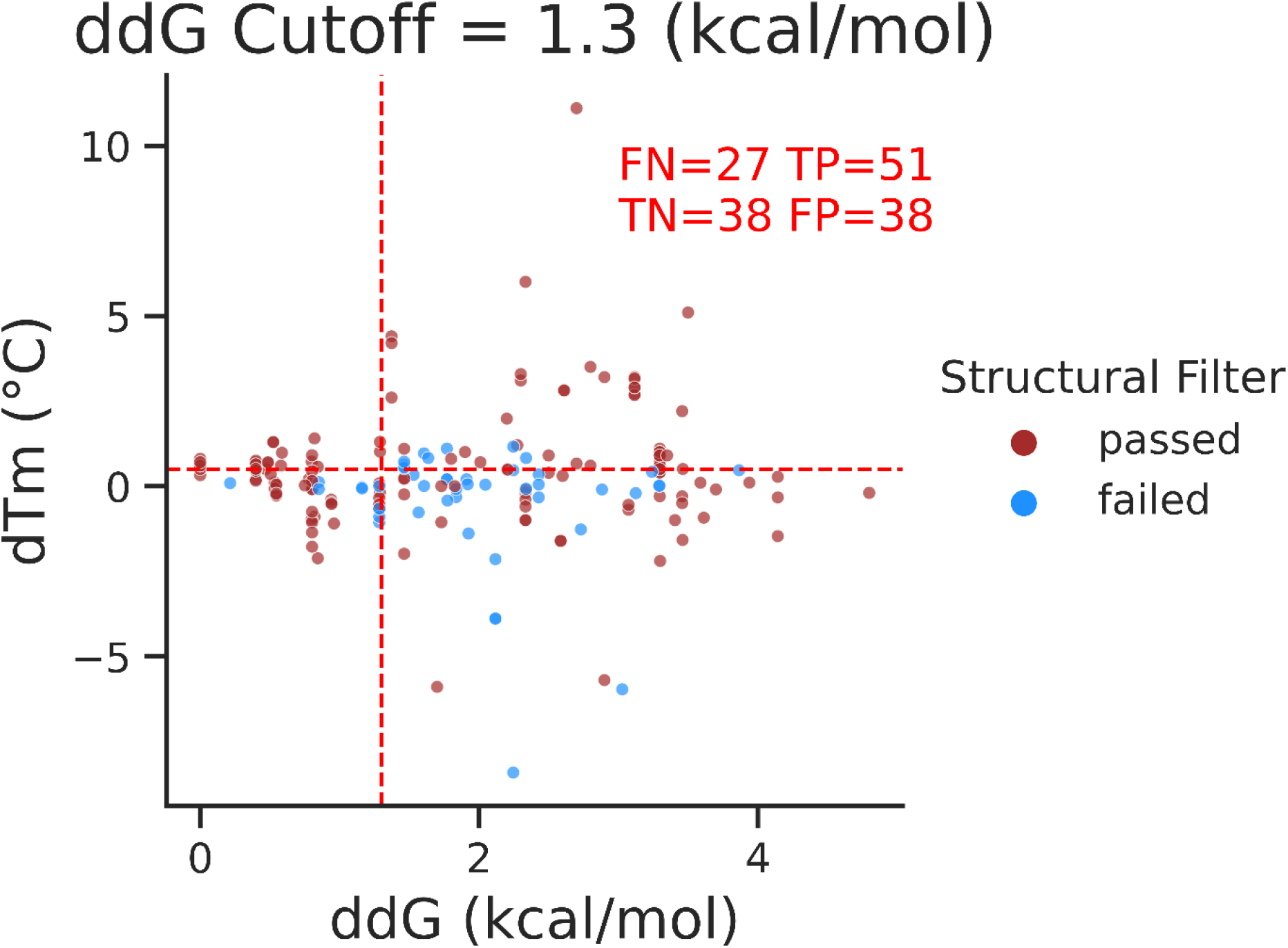
Scatter plot (x-y plot) of dT_m_ VS ddG. X-axis represents the ddG values computed from the consensus sequence method and Y-axis represents the dT_m_ values (difference of experimental T_m_ between the parent and the mutant molecules). The single point mutation pairs are colored based on whether they passed the structural filter or not. The single point mutation pairs which passed the structural filter are classified into TP, FP, TN and FN based on the dT_m_ cutoff of 0.5 °C and the ddG cutoff of 1.3 kcal/mol. The red dotted lines represent the cutoff values for ddG and dT_m_.

**Figure 4:**
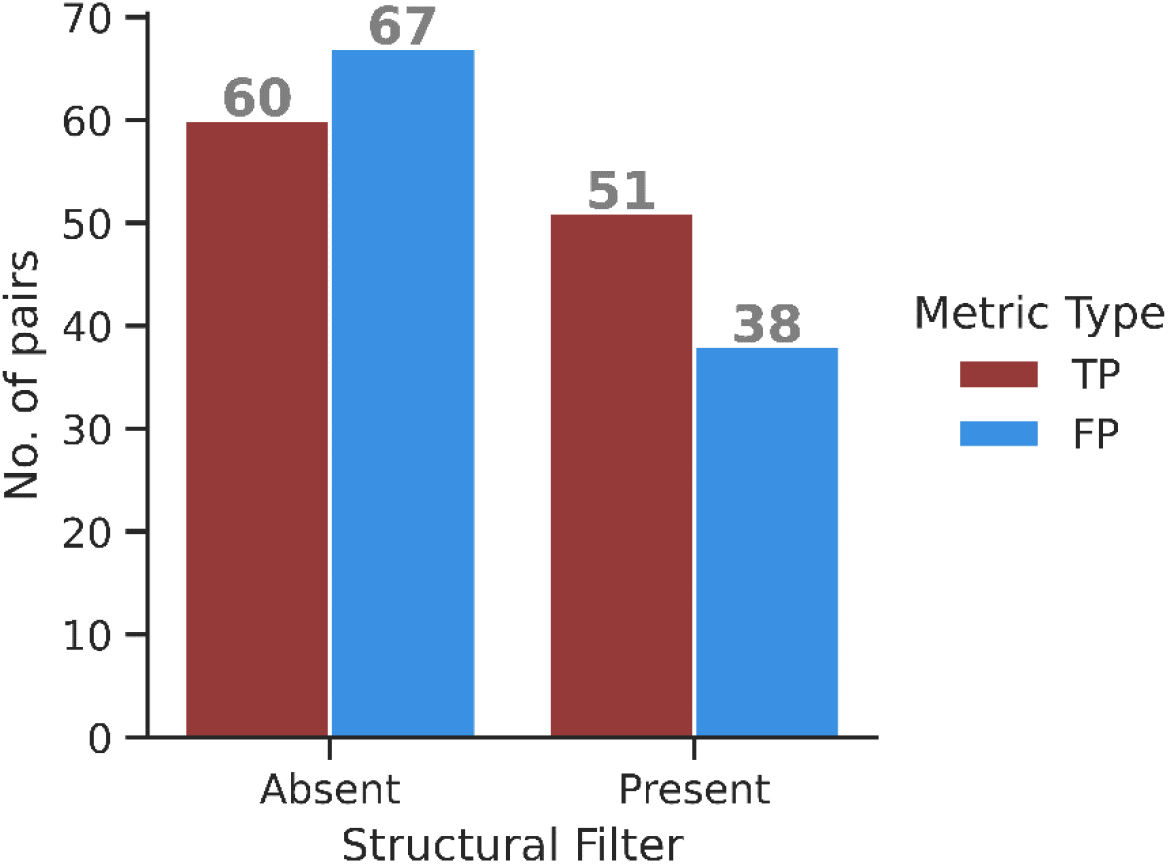
Bar plot showing the effect of structural filtering on reducing FP. TP and FP statistics in this plot are based on the optimal ddG cutoff of 1.3 kcal/mol and dT_m_ cutoff of 0.5 °C.

**Table 4:** Performance comparison between without and with structural filtering. Note: Precision measures the number of correct positive predictions made, it’s defined as TP/(TP+FP). Maximum possible value for precision can be 1.

| Statistics | Without Structural Filtering | With Structural Filtering |
| --- | --- | --- |
| TP | 60 | 51 |
| FP | 67 | 38 |
| TN | 47 | 38 |
| FN | 27 | 27 |
| Precision | 0.47 | 0.57 |

As the available MAbs experimental structures grow, the consensus structure-based MAb close-by residue-pairs can be re-trained with higher confidence. The scoring system can be further improved with consideration of more detailed molecular interaction. Upon reaching certain level of accuracy at which we can use the scoring system to rank molecules’ stability, the method can help to flag any residue pairs in the query molecule that may be liable for stability. This is equivalent to building homology models and manually examining the structures. But our method does not require homology modeling and the manual work, so it’s more systematic and higher throughput.

## Discussion

### Fast stability engineering method in comparison to homology modeling approach

One key advantage of the consensus sequence method in combination with consensus structure-based MAb residue-pair covariance analysis is the fast calculation speed and capability to process a large amount of MAb sequences in high throughput. On average, it takes 19.5 second per MAb molecule. In a typical MAb engineering workflow, the protein engineers may choose homology modeling approach to obtain the same information which can be obtained by residue-pair covariance analysis. However, it takes 3-30 minutes to build a reasonable homology model per MAb molecule. And manually examining the three-dimensional models is even more time consuming. So, the homology modeling approach is not suitable for large scale MAb stability engineering task. Our method is pre-trained by a set of nearly one thousand high resolution crystal structure of MAbs. It takes advantage of conserved structural fold of MAbs to yield the consensus close-by residue pairs. This close-by residue pair information is then used to evaluate the fitness of stability of query MAb based on its sequence. So, our method leverages pre-trained structural information to process only sequence data in order to achieve the goal of fast calculation speed.

### Factors affecting prediction accuracy

The accuracy of consensus sequence prediction and design depends on the following four factors:

1. The gene pool, i.e. the sequence database that is used for multiple sequence alignment (MSA). The larger and more comprehensive of the gene pool, the better chance that the consensus residue can be truly representative. 2. Sequence homology; 3. Sequence count in MSA. 2 and 3 are correlated. The higher the sequence homology, e.g. use germline subset in our application, the fewer of the sequence counts can be used in MSA. Therefore, the sequences selected for MSA are more similar to the query sequence and the consensus sequence yielded from MSA can be more accurate. On the other hand, the fewer number of the sequences in the MSA, the statistical significance would be lower. So, factors 2 and 3 are a tradeoff. The default option for selecting sequences for MSA is germline family, which is a balance between all sequences in IMGT and very few sequences in a given germline. 4. The bias from sequence alignment algorithm used for MSA. For MAb application, several well-developed numbering schemes e.g. IMGT, Chothia, Kabat, and AHo are available. Those numbering systems can generate MSA by using different rules.

### Species specific application

To make the method more accurate, we developed it with certain species consideration. We deployed the method for human MAbs at first. This is our most common use case. All consensus and germline sequences are specific to human MAbs. The consensus method can be further developed for other species given the sequence data is available. Mouse MAb development is the second common use case. We obtained mouse MAb sequences and germline information from IMGT and further developed the method for predicting mouse antibody thermostability.

### Non-antibody applications

The consensus sequence-based stability prediction and design originated from non-antibody applications. The foundation of this method is MSA of a set of sequences with certain level of homology cutoff. In the case of antibody, since the homology of sequences is already very high within the same species, it’s possible to include all sequences from repertoire like IMGT. For higher homology cutoff, germline family and germline subset selections are available. For non-antibody applications, selecting the proper homology cutoff is critical to ensure the validity of MSA and for accurately identifying the consensus residues.

## Conclusion

The goal of our work is to develop a method to help mitigate the thermal stability liability associated with MAbs. We have combined consensus sequence method with consensus structure-based MAb residue-pair covariance analysis to predict thermostabilizing mutations in the query MAb. The theoretical ground of our method is based on the idea that conserved structural fold of MAbs yield consensus close-by residue pairs. This residue pair information is applied to significantly reduce the FP by almost half compared to consensus sequence-based method alone. Major advantages of our method are improved accuracy compared to the consensus sequence method alone and faster computation as well as high-throughput capabilities compared to purely homology modeling based approaches. Future areas of development include enriching the training data for human MAb prediction which will help to further improve accuracy due to the higher statistical significance, developing predictive models for other species’ antibodies, e.g. camelid heavy chain only antibody, and extending applicability to multi-specific antibody engineering.

## Acknowledgement

We would like to acknowledge our colleagues and friends who provided help for our work in this manuscript: Darren Bates, Hannah Catterall, Igor D’angelo, Bram Estes, Fernando Garces, Kevin Graham, Marissa Mock, Austin Rice, Daniel Yoo, and Adam Zalewski.

## References

1. Blankenship K (2020) The top 20 drugs by global sales in 2019. https://www.fiercepharma.com/special-report/top-20-drugs-by-global-sales-2019

2. Le Basle Y, Chennell P, Tokhadze N, et al (2020) Physicochemical Stability of Monoclonal Antibodies: A Review. J. Pharm. Sci. 109:169–190

3. Ewert S, Huber T, Honegger A, Plückthun A (2003) Biophysical properties of human antibody variable domains. J Mol Biol 325:531–553. https://doi.org/10.1016/S0022-2836(02)01237-8

4. McConnell AD, Spasojevich V, Macomber JL, et al (2013) An integrated approach to extreme thermostabilization and affinity maturation of an antibody. Protein Eng Des Sel 26:151–164. https://doi.org/10.1093/protein/gzs090

5. Bondos SE, Bicknell A (2003) Detection and prevention of protein aggregation before, during, and after purification. Anal Biochem 316:223–231. https://doi.org/10.1016/S0003-2697(03)00059-9

6. Weiss IV WF, Young TM, Roberts CJ (2009) Principles, approaches, and challenges for predicting Protein Aggregation Rates and Shelf Life. J. Pharm. Sci. 98:1246–1277

7. Knappik A, Ge L, Honegger A, et al (2000) Fully Synthetic Human Combinatorial Antibody Libraries (HuCAL) Based on Modular Consensus Frameworks and CDRs Randomized with Trinucleotides

8. Seeliger D, Tosatto SCE (2013) Development of Scoring Functions for Antibody Sequence Assessment and Optimization. PLoS One 8:. https://doi.org/10.1371/

9. Couto JR, Christian RB, Peterson JA, Ceriani RL (1995) Designing human consensus antibodies with minimal positional templates. Cancer Res 55:

10. Ewert S, Honegger A, Plückthun A (2004) Stability improvement of antibodies for extracellular and intracellular applications: CDR grafting to stable frameworks and structure-based framework engineering. Methods 34:184–99. https://doi.org/10.1016/j.ymeth.2004.04.007

11. Rouet R, Lowe D, Christ D (2014) Stability engineering of the human antibody repertoire. FEBS Lett 588:269–277. https://doi.org/10.1016/j.febslet.2013.11.029

12. Kulshreshtha S, Chaudhary V, Goswami GK, Mathur N (2016) Computational approaches for predicting mutant protein stability. J Comput Aided Mol Des 30:401–412. https://doi.org/10.1007/s10822-016-9914-3

13. Samish I (2017) Computational Protein Design. Springer New York, New York, NY

14. Steipe B, Schiller B, Plückthun a, Steinbacher S (1994) Sequence statistics reliably predict stabilizing mutations in a protein domain. J. Mol. Biol. 240:188–192

15. Lehmann M, Pasamontes L, Lassen SF, Wyss M (2000) The consensus concept for thermostability engineering of proteins. Biochim Biophys Acta - Protein Struct Mol Enzymol 1543:408–415. https://doi.org/10.1016/S0167-4838(00)00238-7

16. Steipe B (2004) Consensus-based engineering of protein stability: From intrabodies to thermostable enzymes. Methods Enzymol 388:176–186. https://doi.org/10.1016/S0076-6879(04)88016-9

17. Sullivan BJ, Nguyen T, Durani V, et al (2012) Stabilizing proteins from sequence statistics: the interplay of conservation and correlation in triosephosphate isomerase stability. J Mol Biol 420:384–99. https://doi.org/10.1016/j.jmb.2012.04.025

18. Aerts D, Verhaeghe T, Joosten H-JJ, et al (2013) Consensus engineering of sucrose phosphorylase: The outcome reflects the sequence input. Biotechnol Bioeng 110:2563–2572. https://doi.org/10.1002/bit.24940

19. Porebski BT, Buckle AM (2016) Consensus protein design. Protein Eng Des Sel 29:245–51. https://doi.org/10.1093/protein/gzw015

20. Ruiz M, Lefranc MP (2002) IMGT gene identification and Colliers de Perles of human immunoglobulins with known 3D structures. Immunogenetics 53:857–883. https://doi.org/10.1007/s00251-001-0408-6

21. Lefranc MP, Giudicelli V, Duroux P, et al (2015) IMGT R, the international ImMunoGeneTics information system R 25 years on. Nucleic Acids Res 43:D413–D422. https://doi.org/10.1093/nar/gku1056

22. Black SD, Mould DR (1991) Development of hydrophobicity parameters to analyze proteins which bear post- or cotranslational modifications. Anal Biochem 193:72–82

23. Gasser C (2010) Amino Acid Properties. http://www.mcb.ucdavis.edu/courses/bis102/AAProp.html

24. Monsellier E, Bedouelle H (2006) Improving the Stability of an Antibody Variable Fragment by a Combination of Knowledge-based Approaches: Validation and Mechanisms. J Mol Biol 362:580–593. https://doi.org/10.1016/j.jmb.2006.07.044

