## Supporting Tables and Figures for "Improving antibody thermostability based on statistical analysis of sequence and structural consensus data"

**Table S1:** Comparison of the five metrics used for selecting the optimal ddG cutoff. These statistics are based on fixed $dT_m$ cutoff of 0.5 °C. The yellow highlighted row is the selected optimal ddG cutoff.

| ddG Cutoff (kcal/mol) | TP | TN | FP | FN | TP+TN | Precision | Metric 1 | Metric 2 | P Metric | F Metric |
|---|---|---|---|---|---|---|---|---|---|---|
| 0 | 87 | 0 | 114 | 0 | 87 | 0.43 | -869999.24 | -9918 | -9831 | -0.76 |
| 0.1 | 82 | 1 | 113 | 5 | 83 | 0.42 | -81.27 | -9184 | -9102 | -0.72 |
| 0.2 | 82 | 1 | 113 | 5 | 83 | 0.42 | -81.27 | -9184 | -9102 | -0.72 |
| 0.3 | 82 | 2 | 112 | 5 | 84 | 0.42 | -40.27 | -9020 | -8938 | -0.73 |
| 0.4 | 73 | 5 | 109 | 14 | 78 | 0.40 | -13.93 | -7592 | -7519 | -0.65 |
| 0.5 | 70 | 5 | 109 | 17 | 75 | 0.39 | -13.36 | -7280 | -7210 | -0.63 |
| 0.6 | 66 | 13 | 101 | 21 | 79 | 0.40 | -4.42 | -5808 | -5742 | -0.60 |
| 0.7 | 66 | 13 | 101 | 21 | 79 | 0.40 | -4.42 | -5808 | -5742 | -0.60 |
| 0.8 | 66 | 15 | 99 | 21 | 81 | 0.40 | -3.73 | -5544 | -5478 | -0.61 |
| 0.9 | 62 | 31 | 83 | 25 | 93 | 0.43 | -1.25 | -3224 | -3162 | -0.60 |
| 1 | 62 | 35 | 79 | 25 | 97 | 0.44 | -0.99 | -2728 | -2666 | -0.61 |
| 1.1 | 62 | 35 | 79 | 25 | 97 | 0.44 | -0.99 | -2728 | -2666 | -0.61 |
| 1.2 | 62 | 37 | 77 | 25 | 99 | 0.45 | -0.87 | -2480 | -2418 | -0.63 |
| 1.3 | 60 | 47 | 67 | 27 | 107 | 0.47 | -0.38 | -1200 | -1140 | -0.65 |
| 1.4 | 57 | 47 | 67 | 30 | 104 | 0.46 | -0.36 | -1140 | -1083 | -0.50 |
| 1.5 | 52 | 51 | 63 | 35 | 103 | 0.45 | -0.19 | -624 | -572 | -0.08 |
| 1.6 | 52 | 53 | 61 | 35 | 105 | 0.46 | -0.13 | -416 | -364 | 0.13 |
| 1.7 | 50 | 54 | 60 | 37 | 104 | 0.45 | -0.09 | -300 | -250 | 0.67 |
| 1.8 | 49 | 60 | 54 | 38 | 109 | 0.48 | 0.09 | 294 | 343 | -1.83 |
| 1.9 | 48 | 63 | 51 | 39 | 111 | 0.48 | 0.18 | 576 | 624 | -1.25 |
| 2 | 47 | 66 | 48 | 40 | 113 | 0.49 | 0.27 | 846 | 893 | -1.06 |
| 2.1 | 46 | 67 | 47 | 41 | 113 | 0.49 | 0.29 | 920 | 966 | -1.05 |
| 2.2 | 46 | 70 | 44 | 41 | 116 | 0.51 | 0.39 | 1196 | 1242 | -0.92 |
| 2.3 | 42 | 73 | 41 | 45 | 115 | 0.51 | 0.45 | 1344 | 1386 | -0.97 |
| 2.4 | 38 | 80 | 34 | 49 | 118 | 0.53 | 0.64 | 1748 | 1786 | -0.91 |
| 2.5 | 38 | 83 | 31 | 49 | 121 | 0.55 | 0.77 | 1976 | 2014 | -0.87 |
| 2.6 | 37 | 86 | 28 | 50 | 123 | 0.57 | 0.89 | 2146 | 2183 | -0.84 |
| 2.7 | 35 | 87 | 27 | 52 | 122 | 0.56 | 0.89 | 2100 | 2135 | -0.87 |
| 2.8 | 33 | 88 | 26 | 54 | 121 | 0.56 | 0.89 | 2046 | 2079 | -0.89 |
| 2.9 | 31 | 89 | 25 | 56 | 120 | 0.55 | 0.89 | 1984 | 2015 | -0.91 |
| 3 | 30 | 90 | 24 | 57 | 120 | 0.56 | 0.92 | 1980 | 2010 | -0.91 |
| 3.1 | 30 | 93 | 21 | 57 | 123 | 0.59 | 1.11 | 2160 | 2190 | -0.88 |
| 3.2 | 24 | 94 | 20 | 63 | 118 | 0.55 | 0.94 | 1776 | 1800 | -0.95 |
| 3.3 | 4 | 100 | 14 | 83 | 104 | 0.22 | 0.25 | 344 | 348 | -1.12 |
| 3.4 | 3 | 101 | 13 | 84 | 104 | 0.19 | 0.20 | 264 | 267 | -1.11 |
| 3.5 | 1 | 105 | 9 | 86 | 106 | 0.10 | 0.10 | 96 | 97 | -1.08 |

| | | | | | | | | | | |
|---|---|---|---|---|---|---|---|---|---|---|
| 3.6 | 0 | 106 | 8 | 87 | 106 | 0.00 | 0.00 | 0 | 0 | -1.08 |
| 3.7 | 0 | 107 | 7 | 87 | 107 | 0.00 | 0.00 | 0 | 0 | -1.07 |
| 3.8 | 0 | 108 | 6 | 87 | 108 | 0.00 | 0.00 | 0 | 0 | -1.06 |
| 3.9 | 0 | 109 | 5 | 87 | 109 | 0.00 | 0.00 | 0 | 0 | -1.05 |

**Table S2:** Comparison of the five metrics used for selecting the optimal ddG cutoff. These statistics are based on fixed $dT_m$ cutoff of 0.5 °C. The yellow highlighted row is the selected optimal ddG cutoff.

| ddG Cutoff (kcal/mol) | TP | TN | FP | FN | TP+TN | Precision | Metric 1 | Metric 2 | P Metric | F Metric |
|---|---|---|---|---|---|---|---|---|---|---|
| 0 | 78 | 0 | 76 | 0 | 78 | 0.51 | -779998.97 | -5928 | -5850 | -1.03 |
| 0.1 | 73 | 1 | 75 | 5 | 74 | 0.49 | -72.03 | -5402 | -5329 | -0.97 |
| 0.2 | 73 | 1 | 75 | 5 | 74 | 0.49 | -72.03 | -5402 | -5329 | -0.97 |
| 0.3 | 73 | 1 | 75 | 5 | 74 | 0.49 | -72.03 | -5402 | -5329 | -0.97 |
| 0.4 | 64 | 4 | 72 | 14 | 68 | 0.47 | -15.11 | -4352 | -4288 | -0.88 |
| 0.5 | 61 | 4 | 72 | 17 | 65 | 0.46 | -14.40 | -4148 | -4087 | -0.84 |
| 0.6 | 57 | 12 | 64 | 21 | 69 | 0.47 | -3.86 | -2964 | -2907 | -0.87 |
| 0.7 | 57 | 12 | 64 | 21 | 69 | 0.47 | -3.86 | -2964 | -2907 | -0.87 |
| 0.8 | 57 | 14 | 62 | 21 | 71 | 0.48 | -3.15 | -2736 | -2679 | -0.90 |
| 0.9 | 53 | 28 | 48 | 25 | 81 | 0.52 | -0.79 | -1060 | -1007 | -1.25 |
| 1 | 53 | 32 | 44 | 25 | 85 | 0.55 | -0.45 | -636 | -583 | -1.75 |
| 1.1 | 53 | 32 | 44 | 25 | 85 | 0.55 | -0.45 | -636 | -583 | -1.75 |
| 1.2 | 53 | 32 | 44 | 25 | 85 | 0.55 | -0.45 | -636 | -583 | -1.75 |
| 1.3 | 51 | 38 | 38 | 27 | 89 | 0.57 | 0.00 | 0 | 51 | 130000.00 |
| 1.4 | 48 | 38 | 38 | 30 | 86 | 0.56 | 0.00 | 0 | 48 | 100000.00 |
| 1.5 | 47 | 42 | 34 | 31 | 89 | 0.58 | 0.26 | 376 | 423 | 0.63 |
| 1.6 | 47 | 42 | 34 | 31 | 89 | 0.58 | 0.26 | 376 | 423 | 0.63 |
| 1.7 | 47 | 42 | 34 | 31 | 89 | 0.58 | 0.26 | 376 | 423 | 0.63 |
| 1.8 | 47 | 45 | 31 | 31 | 92 | 0.60 | 0.47 | 658 | 705 | 0.14 |
| 1.9 | 46 | 46 | 30 | 32 | 92 | 0.61 | 0.53 | 736 | 782 | 0.00 |
| 2 | 45 | 46 | 30 | 33 | 91 | 0.60 | 0.52 | 720 | 765 | -0.06 |
| 2.1 | 44 | 46 | 30 | 34 | 90 | 0.59 | 0.51 | 704 | 748 | -0.13 |
| 2.2 | 44 | 46 | 30 | 34 | 90 | 0.59 | 0.51 | 704 | 748 | -0.13 |
| 2.3 | 41 | 47 | 29 | 37 | 88 | 0.59 | 0.54 | 738 | 779 | -0.33 |
| 2.4 | 38 | 53 | 23 | 40 | 91 | 0.62 | 0.94 | 1140 | 1178 | -0.50 |
| 2.5 | 38 | 53 | 23 | 40 | 91 | 0.62 | 0.94 | 1140 | 1178 | -0.50 |
| 2.6 | 37 | 56 | 20 | 41 | 93 | 0.65 | 1.19 | 1332 | 1369 | -0.53 |
| 2.7 | 35 | 57 | 19 | 43 | 92 | 0.65 | 1.23 | 1330 | 1365 | -0.58 |
| 2.8 | 33 | 57 | 19 | 45 | 90 | 0.63 | 1.16 | 1254 | 1287 | -0.63 |
| 2.9 | 31 | 57 | 19 | 47 | 88 | 0.62 | 1.09 | 1178 | 1209 | -0.68 |
| 3 | 30 | 58 | 18 | 48 | 88 | 0.63 | 1.15 | 1200 | 1230 | -0.70 |
| 3.1 | 30 | 60 | 16 | 48 | 90 | 0.65 | 1.38 | 1320 | 1350 | -0.68 |
| 3.2 | 24 | 60 | 16 | 54 | 84 | 0.60 | 1.10 | 1056 | 1080 | -0.82 |
| 3.3 | 4 | 63 | 13 | 74 | 67 | 0.24 | 0.24 | 200 | 204 | -1.18 |
| 3.4 | 3 | 64 | 12 | 75 | 67 | 0.20 | 0.20 | 156 | 159 | -1.17 |
| 3.5 | 1 | 68 | 8 | 77 | 69 | 0.11 | 0.11 | 60 | 61 | -1.12 |
| 3.6 | 0 | 69 | 7 | 78 | 69 | 0.00 | 0.00 | 0 | 0 | -1.11 |

| | | | | | | | | | | |
|---|---|---|---|---|---|---|---|---|---|---|
| 3.7 | 0 | 70 | 6 | 78 | 70 | 0.00 | 0.00 | 0 | 0 | -1.09 |
| 3.8 | 0 | 71 | 5 | 78 | 71 | 0.00 | 0.00 | 0 | 0 | -1.08 |
| 3.9 | 0 | 71 | 5 | 78 | 71 | 0.00 | 0.00 | 0 | 0 | -1.08 |

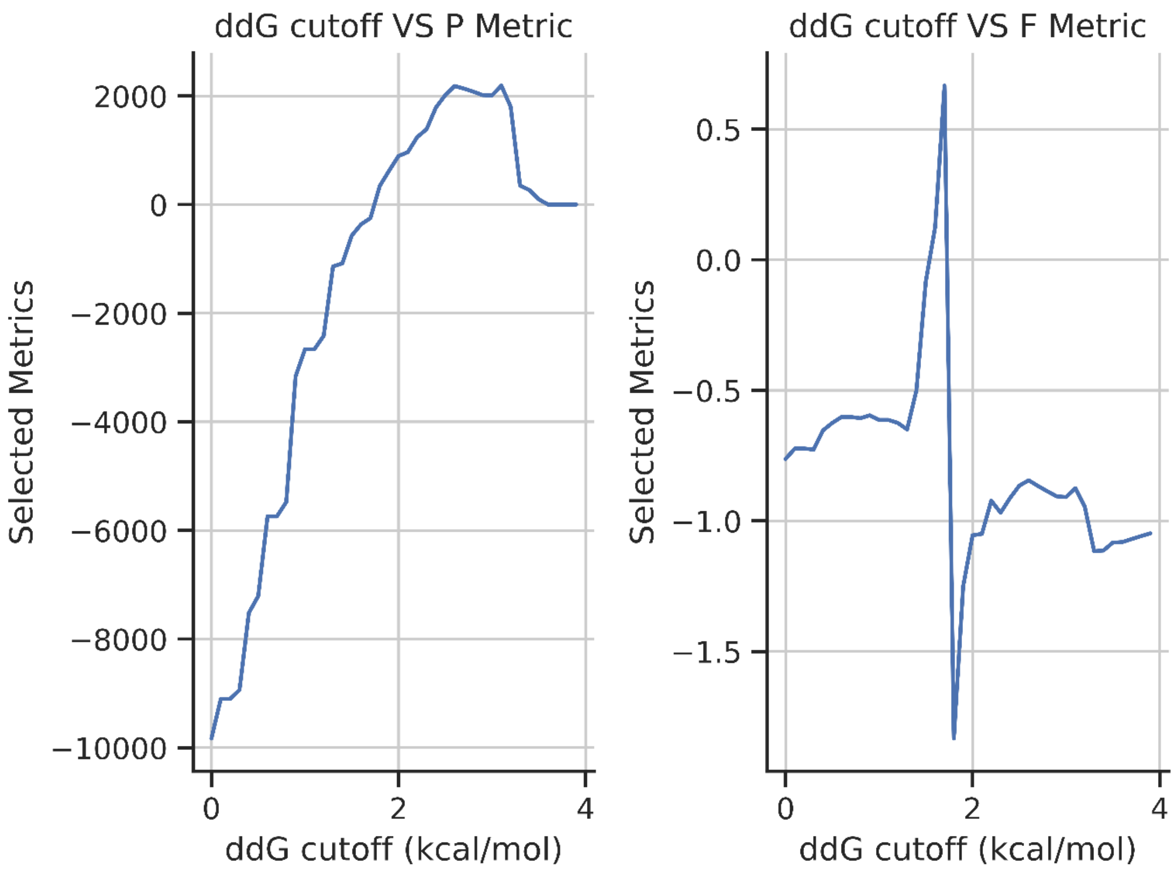


**Figure S1:** ddG cutoff VS P Metric and ddG cutoff VS F Metric are plotted at fixed $dT_m$ cutoff of 0.5 °C. These line plots are showing the values of P Metric and F Metric at different ddG cutoffs. The optimal ddG cutoff using these two metrics is 1.7 kcal/mol.

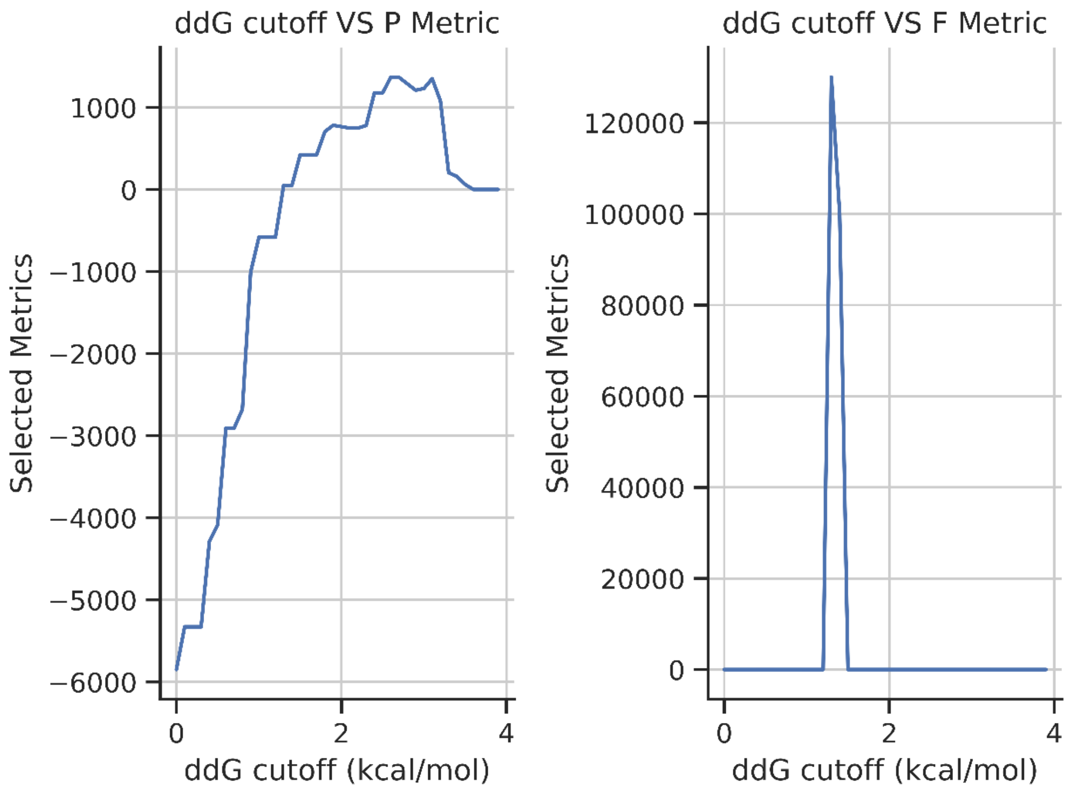


**Figure S2:** ddG cutoff VS P Metric and ddG cutoff VS F Metric are plotted at fixed $dT_m$ cutoff of 0.5 °C. These line plots are showing the values of P Metric and F Metric at different ddG cutoffs. The optimal ddG cutoff using these two metrics is 1.3 kcal/mol.
